# Activation of the neonatal immune system in healthy neonates within hours of birth

**DOI:** 10.1101/2022.02.25.481959

**Authors:** Gaayathri Ariyakumar, Sarah Gee, Abhishek Das, Shraddha Kamdar, Rachel M Tribe, Deena L Gibbons

**Author notes:** These authors have contributed equally to this work and share first authorship (Gaayathri Ariyakumar, Sarah Gee).

## Abstract

It is now established that immune maturation occurs along a defined trajectory in the weeks and months after birth, but the immediate changes that occur within immune cells following birth is less clear. In this study, we monitored the immune profile of neonates via analysis of paired samples (n= 28) of cord blood and heel prick blood taken at varying times post term delivery by caesarean section. This paired approach accounted for the between-subject variability often observed over the first week of life. We identified rapid changes in immune cell populations within hours of birth. Specifically, we observed increased proliferation in effector T cells (but not regulatory T cells) that exhibited an increase in cytokine producing ability and also an increase in the percentage of CD3 T cells over this short time frame. This indicates that the mobilisation of the immune system is immediate post birth, presumably as a response to sudden exposure to the external environment, antigen or stress. Hence, immune development may start to occur more rapidly than previously proposed and as such, to study this trajectory, blood sampling should begin as soon after birth as possible.

## Introduction

Sampling from human neonates, particularly healthy infants born at term, is both difficult and ethically challenging. Much data assessing the neonatal immune system is based on the analysis of cells and cytokines detected in cord blood. Whilst there is evidence that suggests cord blood is not representative of post-natal immunity (1), there are few studies that have assessed whether cord blood can be viewed as indicative of the neonatal immune status directly at birth, and moreover, if any important immune changes occur in first few hours post-delivery. There are myriad immunological challenges as neonates transition from a (relatively) sterile intrauterine environment to a non-sterile extrauterine environment, and thus rapid and well-orchestrated processes are required for neonatal survival. Furthermore, there is substantial between-subject variability, even within the first week of life, which makes understanding changes in immune profiles difficult to interpret on a group basis. However, a greater understanding of the immediate immune changes that are evoked after birth are needed to disentangle why cord blood immune parameters sometimes differ from those in neonatal blood and identify those cells that respond to the immediate challenges post birth and how these responses are manifested. This is particularly pertinent within preterm infants, in whom early onset sepsis, due to acquisition of pathogens transplacentally or via ascending infection during the perinatal period, results in mortality rates as high as 29% (2).

Thus, a greater understanding of the transitional immune events post birth may help in both predicting and assisting neonatal response to infection.

## Materials and Methods

Cord blood and paired heel-prick samples were obtained from 28 neonates born between July 2018 and January 2020 (women provided written informed consent as part of the PROMESA study, REC Approval No. 17/LO/0641). All neonates were born at term gestation by caesarean section (CS). Cord blood was collected at birth and subsequently, paired heel prick blood was obtained within 4 hours of birth (n=18: median 0.4h; range, 0.05 – 3.5h) or between 16 to 48 hours post birth (n=10: median, 22h; range, 17 – 47h) (protocol described in Fig. 1a). Mononuclear cells (MCs) were isolated by Ficoll centrifugation from the different blood samples, washed and then frozen in Cryostor® CS10 (Sigma, UK). Samples were analysed in batches by multiparametric flow cytometry. Cord blood mononuclear cells (CBMCs) were thawed and plated in 96 well round bottom plates (Corning, USA) with or without stimulation with phorbol 12-myristate 13-acetate (PMA) (10 ng/ml) (Sigma), Ionomycin (1 μg/ml) (Sigma), Brefeldin A (20 ng/ml) (Sigma) and Monensin solution (2 μM) (BioLegend). Cells were then stained with one of four antibody panels (Supplementary Table 1) assessing different immune populations and analysed by flow cytometry (4-laser LSR Fortessa, BD). Raw FCS files were analysed using FlowJo (v10.6.2, BD) and the gating strategies are shown in Supplementary Figure 1. Cell populations were excluded from downstream analysis if the event count in the parent population was <30. Analysed flow cytometry populations were imported into an Excel spreadsheet and analysed in R (v4.0.3) to generate principal component analysis plots, boxplots, paired scatter plots, and linear regression correlation plots. Unadjusted p values (*p<0.05; **p<0.01; ***p<0.001, ****p<0.0001) were assessed using two-sided paired Wilcoxon tests (between paired cord blood and heel prick samples).

**Figure 1.**
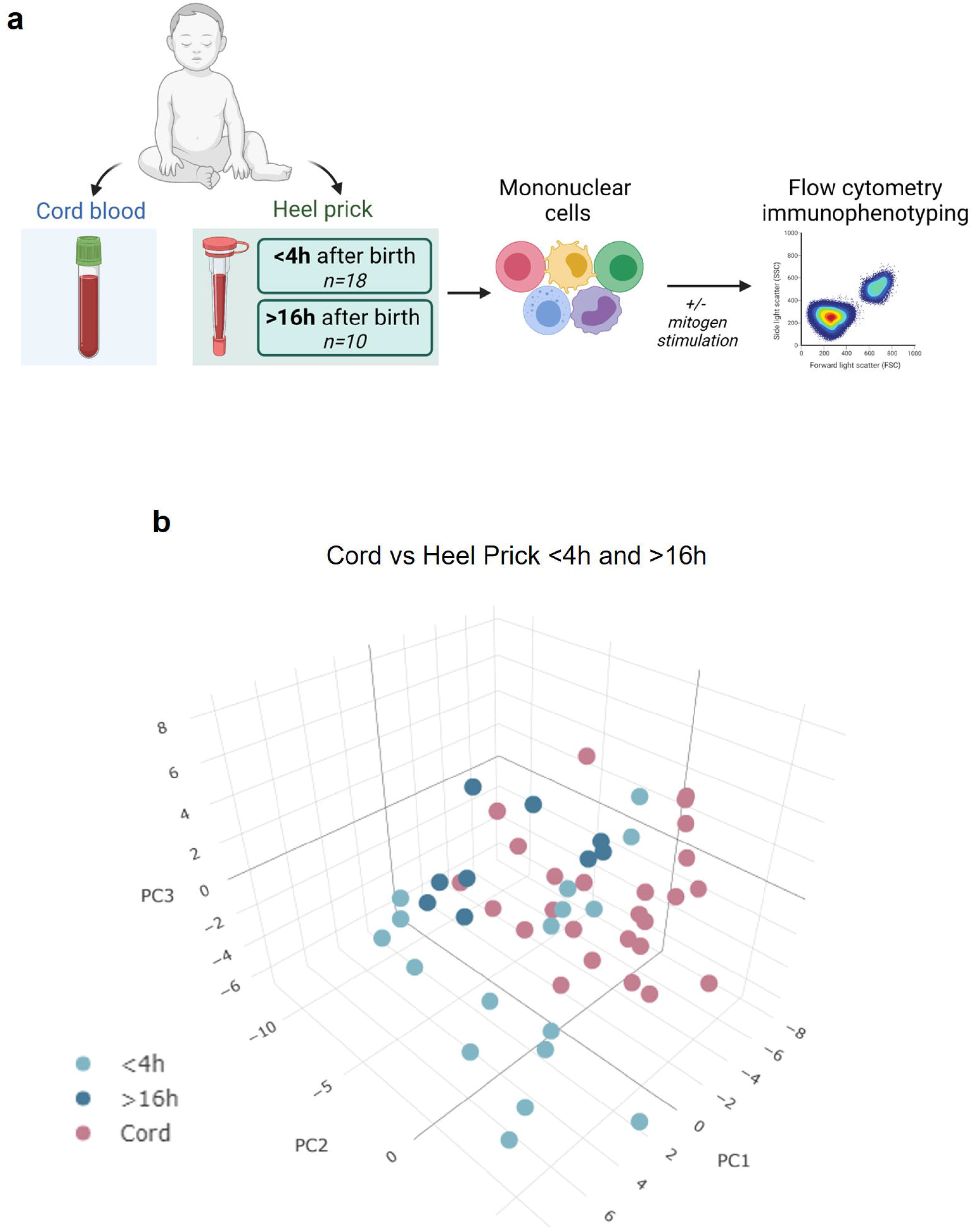
Rapid immune cell changes observed immediately post birth. (a) Experimental design: cord blood and heel prick baseline mononuclear cells were collected at birth or within 48 h post birth. Cells were phenotyped with or without polyclonal stimulation, and cytokine responses and activation markers were measured using multiparametric flow cytometry. Created with BioRender.com (b) 3D PCA visualisation showing the distribution of cord blood samples (pink; n=26) vs paired heel prick baseline samples collected within < 4 h of birth (light blue; n=17) or > 16 h of birth (dark blue; n=9) based on 119 parameters determined by flow cytometry (only samples with complete data set included).

## Results

Cord blood was collected after caesarean section birth and subsequently, paired heel prick blood was obtained within 4 hours of birth (n=18: median 0.4h; range, 0.05 – 3.5h) or between 16 to 48 hours post birth (n=10: median, 22h; range, 17 – 47h) as described in Fig. 1a. Using 3D principal component analysis (PCA) performed on 119 flow cytometry parameters, we compared the immune profile of mononuclear cells isolated from either cord blood or blood samples taken <4h and >16h after birth (Fig 1b). This analysis showed some sample segregation, suggestive of rapid changes in immune cells within the first 48 h of life. Indeed, even those samples taken within 4 h of birth (n=17, light blue, Fig 1b) appeared to segregate from the cord blood samples. When identifying the main drivers of these differences, both cell proliferation (as measured by Ki67 staining) and cytokine producing cells appeared within the top ten. We observed rapid induction of cell proliferation immediately following birth in several CD3^+^ T cell sub-populations. For example, CD4, CD8 and Vδ 1 γ δ T cells exhibited significant elevation in the percentage of Ki67^+^ cells within hours of birth (Fig. 2a-c). This increased proliferation was not associated with cell activation as determined by equivalent CD69^+^ expression on T cells isolated from cord blood or heel pricks (Fig. 2d-f). Indeed, we noted that the proportion of CD3+ T cells increased with time post birth (p<0.001, R=0.35, Fig 2g) consistent with this increased proliferative rate. In comparison, neither Vδ 2 γ δ T cells nor regulatory T cells (Tregs) showed this immediate enhanced proliferation (Fig. 2h,i). In contrast to the proliferation and percentage changes observed in these T cell populations, we did not observe any changes in other populations such as classical monocytes (CD16 CD14) or B cells between the cord blood and heel prick samples (Fig 3 a-b), and there was no correlation with B cells and time post birth (R=0.0007, Fig 3c).

**Figure 2.**
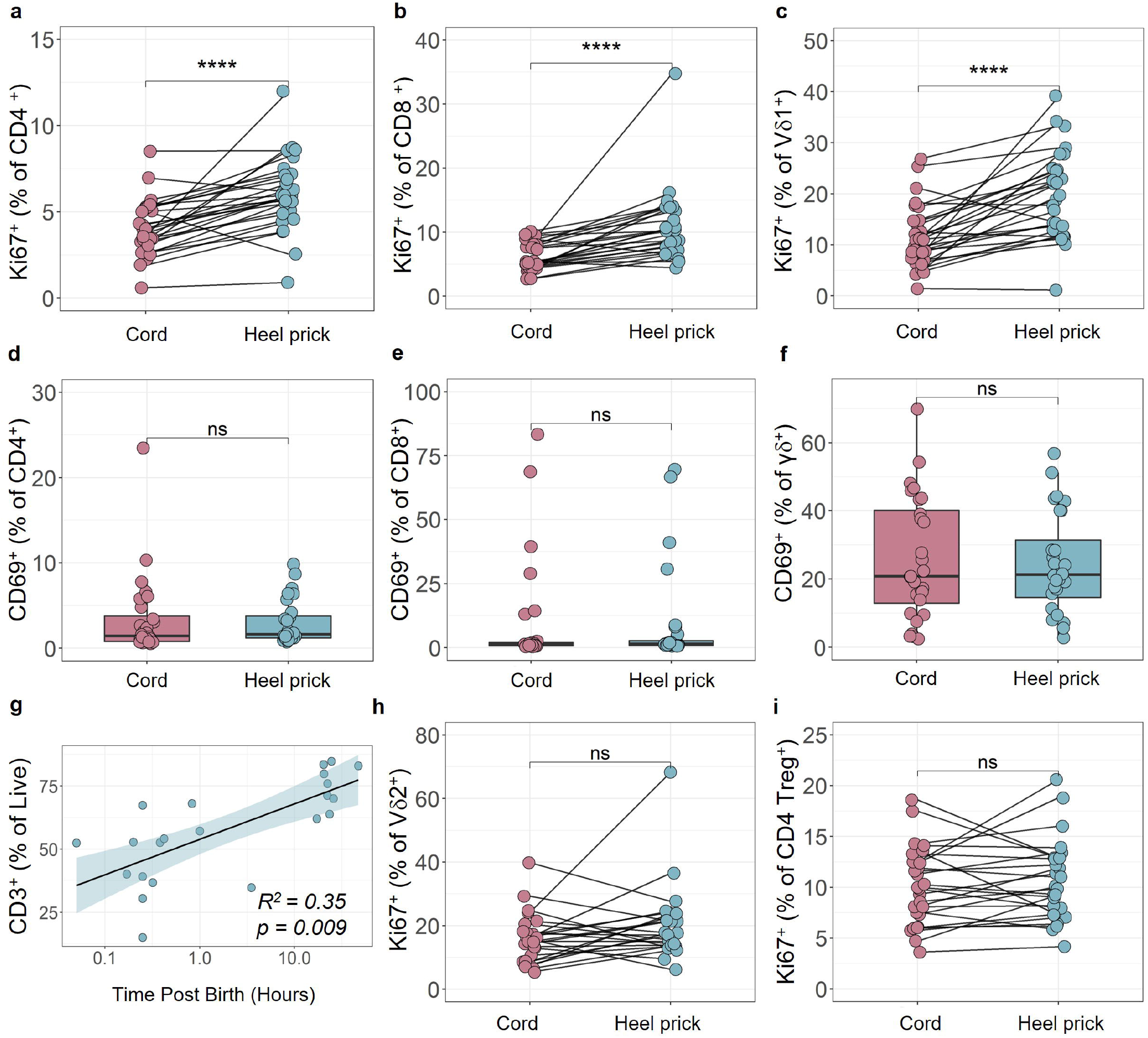
Enhanced proliferation in neonatal T cells observed immediately post birth. (a-c) The percentage of proliferating (Ki67^+^) CD4^+^ (a), CD8^+^ (b), and Vδ1+ γ δ (c) cells from paired cord blood and heel prick baseline samples (n=28). (d-f) The percentage of activated (CD69^+^;) CD4^+^ (d), CD8^+^ (e) and γδ ^+^ (f) T cells from paired cord blood and heel prick baseline samples (n=28). (g) Time dependent linear regression analysis of the percentage of live CD3^+^ T cells from heel prick baseline blood samples collected within 48 h post birth (n=28). (h-i) The percentage of proliferating (Ki67^+^) Vδ2^+^ γδ (h), and CD4^+^ Treg (i) cells. Box and whisker plots follow standard Tukey representations and indicate the median (central line), the interquartile range (upper line = 75^th^ percentile, lower line = 25^th^ percentile), and the whiskers represent 1.5 × 75^th^ /25^th^ percentile. P values (NS, non-significant; ****p<0.0001) for all paired data between cord and heel prick baseline samples were assessed using two-sided paired Wilcoxon tests.

**Figure 3.**
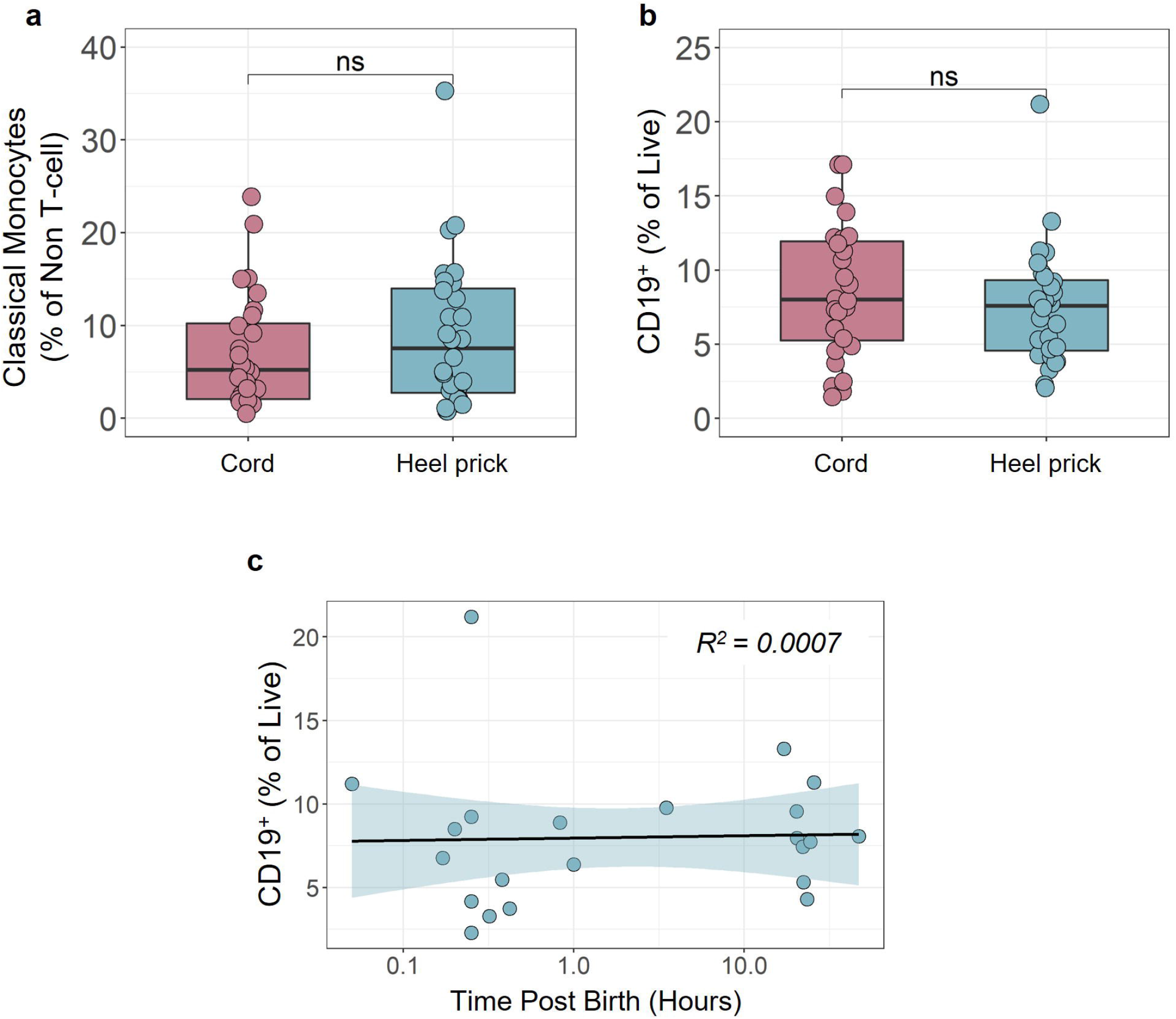
Percentage of neonatal classical monocytes and B cells remain unchanged immediately post birth. Proportion of (a) classical monocytes (CMCs) and (b) CD19^+^ B cells was measured by multiparametric flow cytometry in cord blood and paired heel prick baseline samples collected within 48 h post birth (n=28). (c) Time dependent linear regression analysis of CD19^+^ B-cells from heel prick baseline blood samples collected within 48 h post birth (n=28). Box and whisker plots follow standard Tukey representations and indicate the median (central line), the interquartile range (upper line = 75^th^ percentile, lower line = 25^th^ percentile), and the whiskers represent 1.5 × 75^th^ /25^th^ percentile. NS, non-significantfor all paired data between cord and heel prick baseline samples were assessed using two-sided paired Wilcoxon tests.

Consistent with a potential mobilisation of the neonatal immune system immediately post birth, we also observed a rapid (< 4 h) increase in the percentage of CD4, CD8, γ δ T cells and NK cells producing TNF (Fig 4a-e). Interestingly, this was not as apparent in cells isolated from blood > 16 h post birth (Fig 4a-e) suggesting a rapid, transitory effect. Consistent with the observed increased proliferation, the percentage of IL-2 producing CD4, CD8, γ δ T cells and NK cells was also significantly increased in the samples taken within hours of birth (Fig 4f-j).

**Figure 4.**
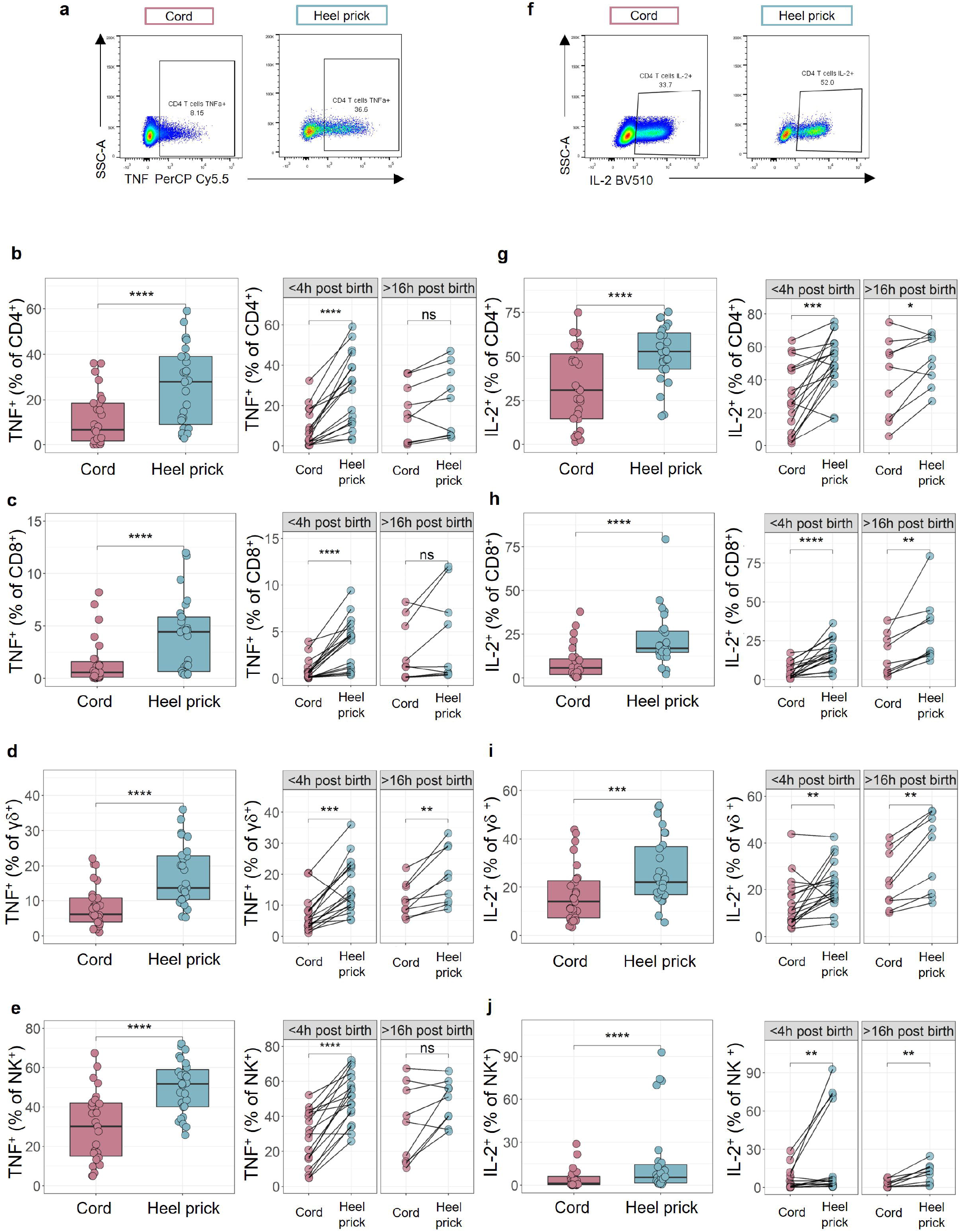
Elevated percentages of cytokine-expressing immune cells observed immediately post birth. Cytokine production was measured by multiparametric flow cytometry in mononuclear cells isolated from cord blood or paired heel prick baseline samples in response to polyclonal stimulation [PMA (10 ng/ml), ionomycin (1 μg/ml), brefeldin A (20 ng/ml) and monensin solution (2 μM) at 37 C for 4 h]. (a) Representative flow plots of TNF in CD4^+^ T cells from cord and heel prick baseline samples. (b-e) The percentage of TNF expressing cells in CD4^+^ (b), CD8^+^ (c) and γδ ^+^ (d) and NK (e) cells from cord blood (n=28) compared to heel prick baseline samples collected <4 h (n=18) or >16 h post birth (n=10). (f) Representative flow plots of IL-2 in CD4^+^ T cells from cord and heel prick baseline samples. (g-j) The percentage of IL-2 expressing cells in CD4^+^ (g), CD8^+^ (h) and γδ ^+^ (i) and Natural Killer (NK) (j) cells from cord blood (n=28) compared to heel prick baseline samples collected <4 h (n=18) or >16 h post birth (n=10). Box and whisker plots follow standard Tukey representations and indicate the median (central line), the interquartile range (upper line = 75^th^ percentile, lower line = 25^th^ percentile), and the whiskers represent 1.5 × 75^th^ /25^th^ percentile. P values (***p<0.001, ****p<0.0001) for all paired data between cord and heel prick baseline samples were assessed using two-sided paired Wilcoxon tests.

## Discussion

At birth, the transition from *in utero* to an external environment presents a need for rapid physiological adaptation coupled with an ability to respond to microbial challenges. Whilst it is known that there are conspicuous developmental changes within the first few weeks of life along a very defined trajectory (3) (1) (4), the rapid immune changes we observed in healthy neonates within a few hours of birth have not previously been anticipated. An earlier study reported only 12 differentially expressed genes in transcriptomic analysis of whole blood between day 0 and day 1 although extensive changes thereafter (3). Although some phenotypic cellular changes were also observed in that study, most were reductions (or no change) in normalised cell numbers over the first week of life. Indeed, in our study, despite observing some changes in the percentages of CD3^+^ T cells in this very early time window, the majority of cell types were also unchanged, and most alterations observed were related to the functional state of the different cell types, as opposed to a proportional change in the numbers of different cell populations. Plasma concentrations of some proteins, such as complement and acute phase proteins are known to increase rapidly post birth with some (e.g., serum amyloid A), showing just a transient rise indicative of an acute response (5). Indeed, much of the changes in both proliferation and cytokine potential observed were not as apparent in samples taken >16 h post birth, suggesting our observations are a transient response to myriad challenges immediately post birth. It is known that several cell types, as well as cytokines, are altered during the natural birth process (6) (7) (8) (9). Our samples, however, were all taken after planned caesarean deliveries, and as such, would not have been subjected to the magnitude of hormonal surges or stress response associated with labour and vaginal delivery (10). It should also be noted that maternal immune cells do not differ pre and post caesarean section (11), suggesting we are observing a neonatal specific response to the extrauterine environment. Similarly, whilst proliferating T cells have been observed in neonates born to mothers with chorioamnionitis (4), all neonates in this study were born to healthy mothers with no evidence or record of exposure to placental inflammation or infection.

It is intriguing, therefore, to consider what may be driving this rapid mobilisation of the immune system. Without a comparison to vaginally delivered infants, it is impossible to be sure that the immediate proliferative response is not merely related to the recovery from any anaesthetic used during surgery. Indeed, CS delivered neonates show a heightened behavioural response to stress assessed 2 h after birth (12). Assuming the observed responses are not related solely to mode of delivery, it could be possible that this is this an antigen specific response or a more general response to stress such as cold shock or changes in oxygen tension immediately upon delivery. Environmental temperature has been shown to affect immune parameters in neonatal pigs; where piglets maintained in a cold environment of 18 C and challenged with LPS had greater serum concentrations of TNF and cortisol compared to pigs maintained at 34 C (13). Thus, enhanced TNF as observed, could be related to the cold shock of delivery.

Similarly, could these responses be related to microbial/commensal exposures post birth? Neonatal microbial colonisation begins immediately where the gut microbiotas are highly dynamic and individualised during the neonatal period. This is dependent both on mode of delivery, driven by skin contact of the baby with the mother, and breast feeding (14-16). Also, the disrupted transmission of the maternal gastrointestinal bacteria observed in C-section babies associates with a substantially higher relative abundance of opportunistic pathogens commonly associated with the healthcare and hospital environment (17) which may potentially drive greater responses. This exposure to microbial antigens facilitates the development and maturation of the neonatal immune system (18). Hence, this initial rapid colonisation may be at least initiating the responses observed.

The increase in IL-2-producing cells would be consistent with the enhanced proliferation of T cells for which IL-2 is an autocrine growth factor, induced downstream of T cell activation. This notwithstanding, we did not observe any enhanced proliferation in Tregs which are usually more responsive to IL-2 stimulation. Indeed, the specific induction of effector (but not regulatory) T cell proliferation may facilitate an immediate effector T cell response to multiple antigen exposure without inhibition. This study did not assess T cell clonality, however, and hence we could not establish whether this is a specific T cell receptor mediated response to antigen. Whilst the increase in proliferation observed was rapid, it must be noted that Ki67 expression, a marker extensively used to indicate cell proliferation, only delineates cells that are not in G0 (19) and how long these cells remain in, or indeed if they exit G1, cannot be elucidated. Similarly, not all cells showed an increase in Ki67 immediately post birth. Notably, the percentage of B cells remained constant, despite these cell populations being known to expand at least in the first few months post birth (4).

Our study suggests that the neonatal immune system rapidly changes within the first few hours of birth as the neonate transitions from the protected *in utero* environment to the diverse microbial environment. This has broad implications for a better understanding of how the neonate protects itself in these early hours/days and how this could be enhanced therapeutically. Our study highlights a greater need to focus on this very early development state of the human neonatal immune system.

## Acknowledgements

We thank the mothers and their neonates for blood collection and all the midwives at STH for sample collection. We would also like to acknowledge Evolve Biosystems for funding (RT and DG) towards the neonatal blood sample collection and the staff involved (Niamh Kelly, Lucy McMillan, Sarah Kheirallah and Jiadai Mi). GA is supported by Action Medical Research grant held by DG (GN2790); SG is supported by a MRC-KCL Doctoral Training Partnership in Biomedical Sciences (MR/N013700/1), RMT by Tommy’s (Charity No. 1060508) and Borne (1167073). The research waslllalso supported by the National Institute for Health Research (NIHR) Biomedical Research Centre based at Guy’s and St Thomas’ NHS Foundation Trust (GSTT) and King’s College London (KCL) (part of the King’s Health Partners Academic Sciences Centre). The views expressed are those of the authors and not necessarily those of the NHS, the NIHR or the Department of Health.

## Author Contributions

GA: Flow cytometry analysis and manuscript drafting; SG, SK: Sample processing, flow cytometry and analysis; SK, AD: panel design, flow cytometry. DG and RT: Study conception, experimental design and manuscript drafting. All authors reviewed drafts of the manuscript prior to submission.

## References

1. Olin A, Henckel E, Chen Y, Lakshmikanth T, Pou C, Mikes J, et al. Stereotypic Immune System Development in Newborn Children. Cell. 2018;174(5):1277–92 e14.

2. Stoll BJ, Puopolo KM, Hansen NI, Sanchez PJ, Bell EF, Carlo WA, et al. Early-Onset Neonatal Sepsis 2015 to 2017, the Rise of Escherichia coli, and the Need for Novel Prevention Strategies. JAMA Pediatr. 2020;174(7):e200593.

3. Lee AH, Shannon CP, Amenyogbe N, Bennike TB, Diray-Arce J, Idoko OT, et al. Dynamic molecular changes during the first week of human life follow a robust developmental trajectory. Nat Commun. 2019;10(1):1092.

4. Kamdar S, Hutchinson R, Laing A, Stacey F, Ansbro K, Millar MR, et al. Perinatal inflammation influences but does not arrest rapid immune development in preterm babies. Nat Commun. 2020;11(1):1284.

5. Bennike TB, Fatou B, Angelidou A, Diray-Arce J, Falsafi R, Ford R, et al. Preparing for Life: Plasma Proteome Changes and Immune System Development During the First Week of Human Life. Front Immunol. 2020;11:578505.

6. Tornblom SA, Maul H, Klimaviciute A, Garfield RE, Bystrom B, Malmstrom A, et al. mRNA expression and localization of bNOS, eNOS and iNOS in human cervix at preterm and term labour. Reprod Biol Endocrinol. 2005;3:33.

7. Garcia-Serna AM, Martin-Orozco E, Hernandez-Caselles T, Morales E. Prenatal and Perinatal Environmental Influences Shaping the Neonatal Immune System: A Focus on Asthma and Allergy Origins. Int J Environ Res Public Health. 2021;18(8).

8. Nepsha OS, Nikitina IV, Donnikov AE, Bystritsky AA, Mullabaeva SM, Pavlovich SV, et al. Comparison of the Expression Profiles of Immune Response Gene mRNA in Umbilical and Venous Blood of Newborns of the First Day of Life. Bull Exp Biol Med. 2016;160(6):791–4.

9. Werlang ICR, Mueller NT, Pizoni A, Wisintainer H, Matte U, Costa S, et al. Associations of birth mode with cord blood cytokines, white blood cells, and newborn intestinal bifidobacteria. PloS one. 2018;13(11):e0205962.

10. Tribe RM, Taylor PD, Kelly NM, Rees D, Sandall J, Kennedy HP. Parturition and the perinatal period: can mode of delivery impact on the future health of the neonate? J Physiol. 2018;596(23):5709–22.

11. van Egmond A, van der Keur C, van Beelen E, Scherjon SA, Claas FHJ. No direct effect of an elective caesarean section on the phenotypic and functional characteristics of maternal peripheral blood T lymphocytes. Hum Immunol. 2016;77(10):898–904.

12. Chis A, Vulturar R, Andreica S, Prodan A, Miu AC. Behavioral and cortisol responses to stress in newborn infants: Effects of mode of delivery. Psychoneuroendocrinology. 2017;86:203–8.

13. Carroll JA, Burdick NC, Chase CC, Jr., Coleman SW, Spiers DE. Influence of environmental temperature on the physiological, endocrine, and immune responses in livestock exposed to a provocative immune challenge. Domest Anim Endocrinol. 2012;43(2):146–53.

14. Stewart CJ, Ajami NJ, O’Brien JL, Hutchinson DS, Smith DP, Wong MC, et al. Temporal development of the gut microbiome in early childhood from the TEDDY study. Nature. 2018;562(7728):583–8.

15. Heikkila MP, Saris PE. Inhibition of Staphylococcus aureus by the commensal bacteria of human milk. J Appl Microbiol. 2003;95(3):471–8.

16. Dominguez-Bello MG, Costello EK, Contreras M, Magris M, Hidalgo G, Fierer N, et al. Delivery mode shapes the acquisition and structure of the initial microbiota across multiple body habitats in newborns. Proc Natl Acad Sci U S A. 2010;107(26):11971–5.

17. Shao Y, Forster SC, Tsaliki E, Vervier K, Strang A, Simpson N, et al. Stunted microbiota and opportunistic pathogen colonization in caesarean-section birth. Nature. 2019;574(7776):117–21.

18. Simon AK, Hollander GA, McMichael A. Evolution of the immune system in humans from infancy to old age. Proc Biol Sci. 2015;282(1821):20143085.

19. Di Rosa F, Cossarizza A, Hayday AC. To Ki or Not to Ki: Re-Evaluating the Use and Potentials of Ki-67 for T Cell Analysis. Front Immunol. 2021;12:653974.

