## Supplementary figures and images for "Activation of the neonatal immune system in healthy neonates within hours of birth"

### Supplementary Figure 1

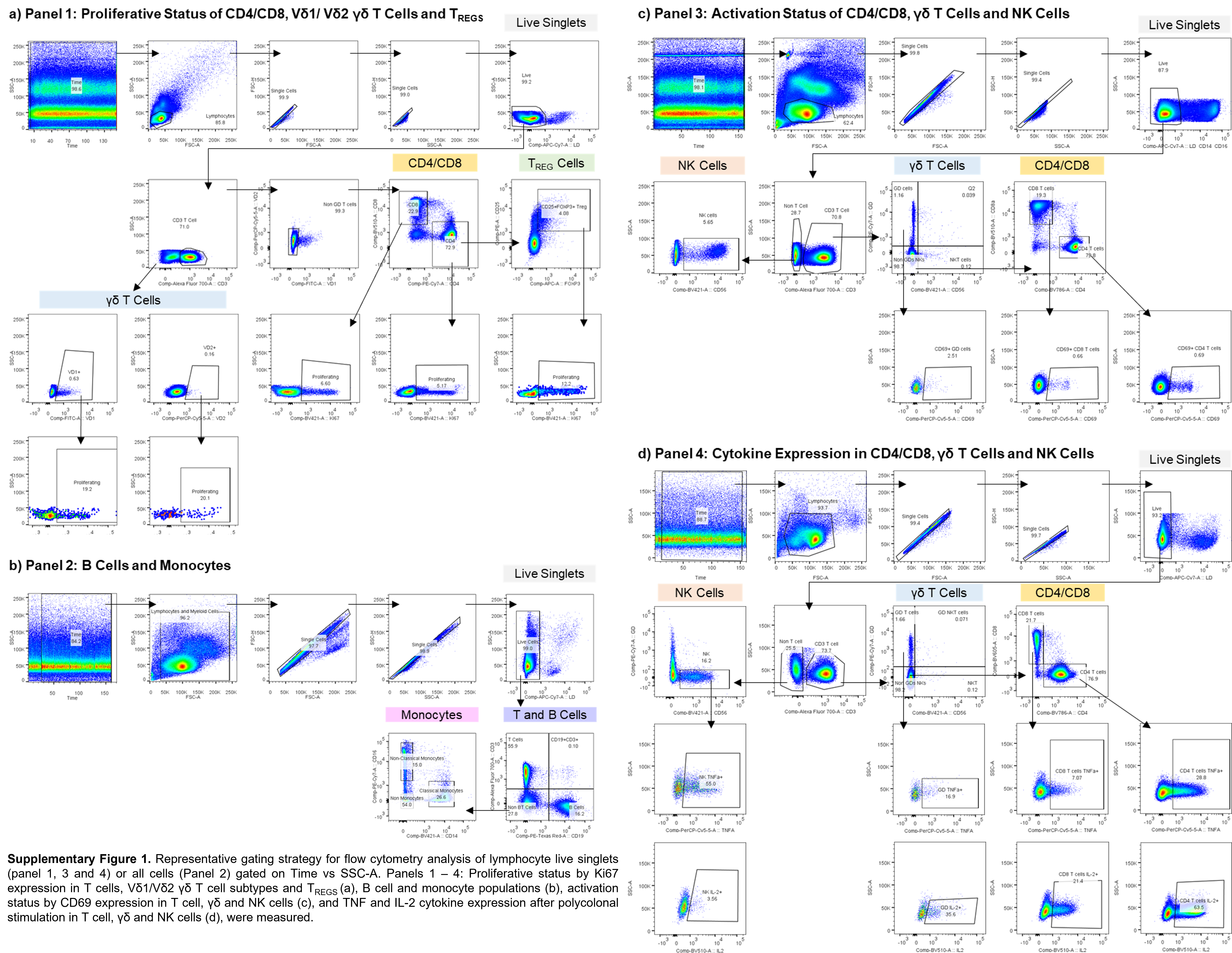

### Supplementary Table 1

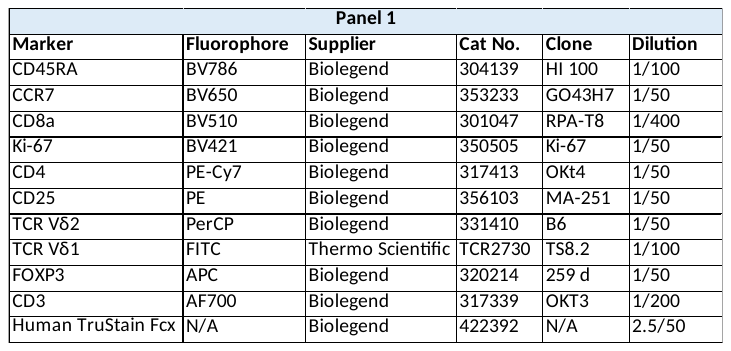

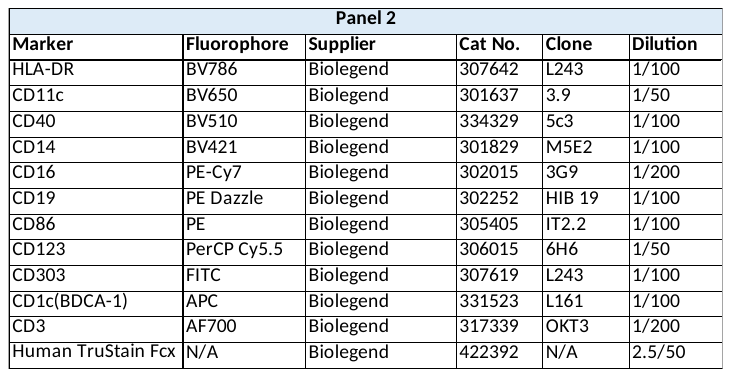

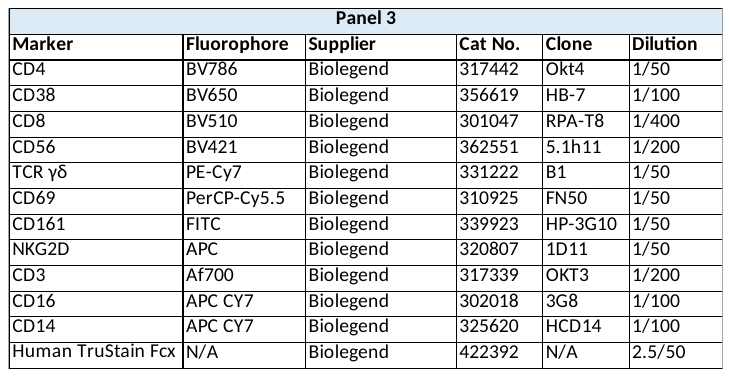

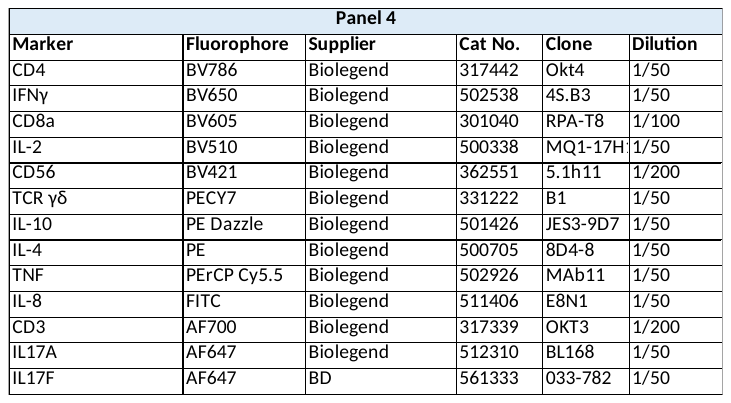
